# Limb bone scaling in hopping macropods and quadrupedal artiodactyls

**DOI:** 10.1101/256768

**Authors:** Michael Doube, Alessandro A Felder, Melissa Y Chua, Kalyani Lodhia, Michał M Kłosowski, John R Hutchinson, Sandra J Shefelbine

**Affiliations:** Department of Bioengineering, Imperial College London, London SW7 2AZ, UK; Skeletal Biology Group, The Royal Veterinary College, Royal College Street, London NW1 0TU, UK; Structure and Motion Laboratory, The Royal Veterinary College, North Mymms, Hatfield, Hertfordshire AL9 7TA, UK; Department of Mechanical & Industrial Engineering, Northeastern University, 334 Snell Engineering Center, 360 Huntington Avenue, Boston, MA 02115, USA

## Abstract

Bone adaptation is modulated by the timing, direction, rate, and magnitude of mechanical loads. To investigate whether frequent slow, or infrequent fast, gaits could dominate bone adaptation to load, we compared scaling of the limb bones from two mammalian herbivore clades that use radically different high-speed gaits, bipedal hopping (suborder Macropodiformes; kangaroos and kin) and quadrupedal galloping (order Artiodactyla; goats, deer and kin). Forelimb and hindlimb bones were collected from 20 artiodactyl and 15 macropod species (body mass *M* 1.05 – 1536 kg) and scanned in computed tomography or X-ray microtomography. Second moment of area (*I*_max_) and bone length (*l*) were measured. Scaling relations (*y* = *ax^b^*) were calculated for *l* vs *M* for each bone and for *I*_max_ vs *M* and *I*_max_ vs *l* for every 5% of length. *I*_max_ vs *M* scaling relationships were broadly similar between clades despite the macropod forelimb being nearly unloaded, and the hindlimb highly loaded, during bipedal hopping. *I*_max_ vs *l* and *l* vs *M* scaling were related to locomotor and behavioural specialisations. Low-intensity loads may be sufficient to maintain bone mass across a wide range of species. Occasional high-intensity gaits might not break through the load sensitivity saturation engendered by frequent low-intensity gaits.

## Introduction

During daily rest and activity in development, growth, and adulthood, bones experience a range of mechanical loading conditions that relate to each behaviour’s physical intensity. Kangaroos, wallabies and their macropodiform kin are famed for their hopping hindlimb gait which they use for bursts of efficient high-speed locomotion [1–3]. They are less well known for their slower pentapedal gait, wherein their powerful tail acts as the third point of a tripod with the forelimbs during hindlimb protraction [4] (Fig. 1). The pentapedal gait is used during grazing, and along with other slow-speed activities, dominates macropods’ locomotor behaviour. Tammar wallabies *(Macropus eugenii)* spend up to twice as much time per day in pentapedal walking than in bipedal hopping (6% vs 3-5%), and both gaits are eclipsed in the locomotor time budget by bipedal standing (50-70%), quadrupedal crouching (15-30%), and bipedal rearing (3-12%) [3,5,6]. During hopping, the forelimbs are held away from ground contact for the entire stride cycle and thus are relatively unloaded [2], while hindlimb tissues experience near-ultimate stresses from ground reaction forces and muscle-tendon action, especially in larger Macropodiformes [7]. The tail’s role in pentapedal locomotion during slow-speed locomotion might enable reduced forelimb mass, potentially assisting more efficient bipedal hopping [4]. In extinct sthenurine macropods, the thoracic limb displays features of a browsing adaptation with elongated manus, reduced lateral digits, slender radius, ulna and humerus, and a ‘human-like’ scapula, which may have enabled these animals to forage browse above their heads [8]. Hopping is likely not possible at body mass over ~160 kg, at which the distal tendons’ safety factor (ratio of actual to ultimate stress) drops below 1, meaning that extinct ‘giant kangaroos’ would have used slower gaits [7–10].

**Figure 1.**
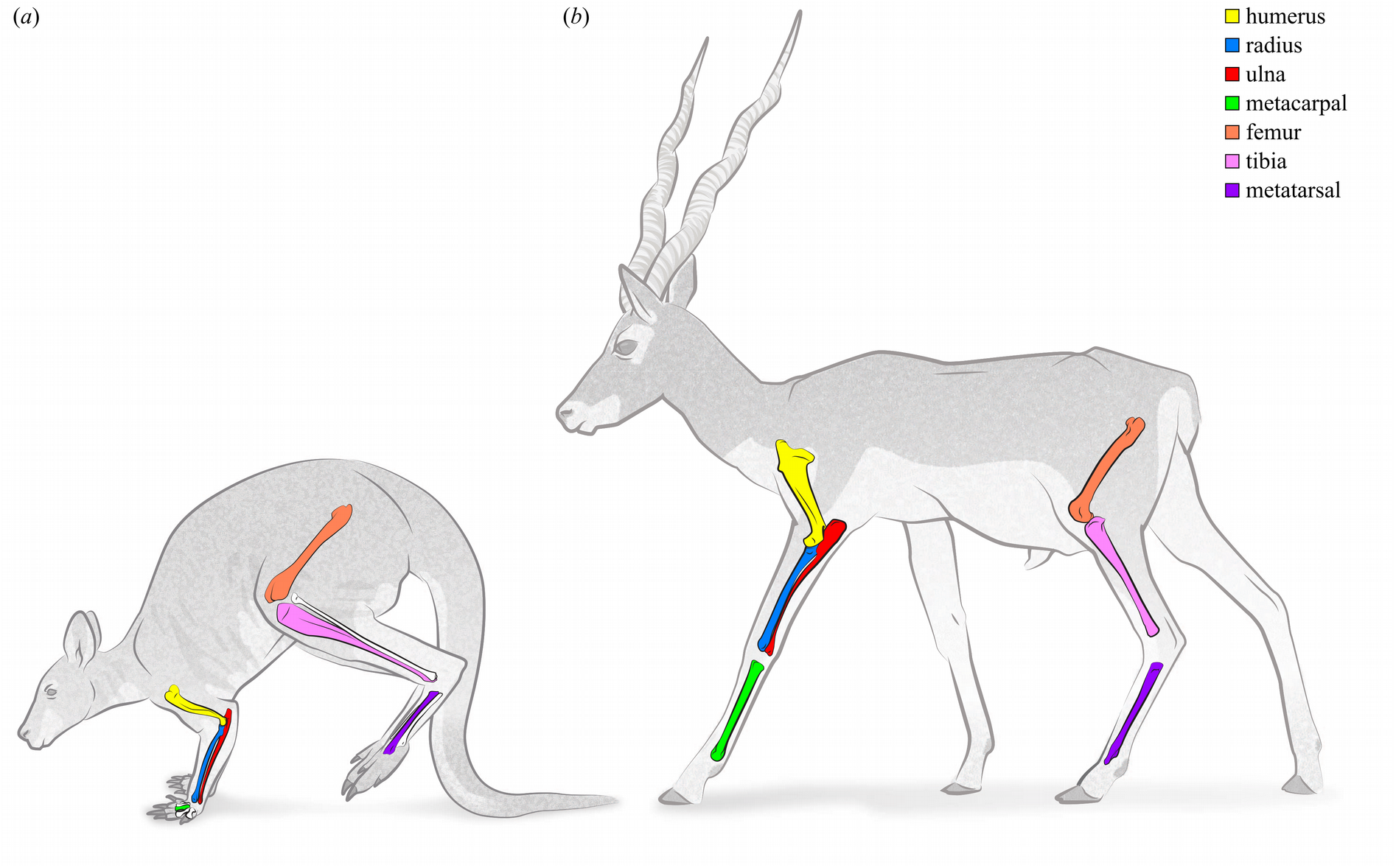
Bennett’s wallaby *(Macropus rufogriseus)* in the hindlimb suspension phase of the pentapedal gait (a) and blackbuck *(Antelope cervicapra)* in a lateral sequence walk (b) indicating the limb bones measured in the study. These two species have femora of similar length (199 mm and 186 mm respectively) and are presented here approximately to scale. Drawing by Manuela Bertoni may be reused under the CC BY licence.

In contrast to macropods, artiodactyl mammals (even-toed ungulates in the eutherian lineage; deer, sheep, camels and kin) have limited manual dexterity, and quadrupedal gaits in which the loads are spread evenly among fore- and hindlimbs during both slow and fast gaits, reflected in similarity of forelimb and hindlimb bones’ cross-sectional properties [11]. Artiodactyls and macropods spend a large proportion of their time grazing or resting as they are foregut fermenter herbivores [12] and may be considered ecological equivalents [13]. Extinct giant macropods could not hop due to tissue strength being exceeded by scaling of muscle and tendon stress [9], yet even the largest artiodactyls retain high-speed gaits. Bison, buffalo, and giraffe are capable of galloping [14, 15], while hippopotami achieve high land speeds by a fast walk or trot [16]. Scaling of limb bones in artiodactyls is relatively well characterised, exhibiting isometric or modestly allometric patterns across their 3-order of magnitude body mass range [17–20].

The distributions of occasional maximal loads and habitual moderate loads vary within the skeleton and depend on locomotor activity, which should appear as a morphological signal in clades that adopt very different characteristic gaits [21]. Although direct bone strain gauge data do not exist, positive allometry of hindlimb muscle physiological cross-sectional area, reduced duty factor with increasing speed, and constant effective mechanical advantage of hindlimb joints, likely lead to relatively increased muscle force, and increased stress and reduced safety factors in larger macropods’ hind limb bones [10,22,23]. Bennett (2000) pointed out that kangaroos’ tibial cross-sections (section modulus *Z* and second moment of area *I,* which relate to fracture strength and resistance to bending respectively) scale more strongly than other quadrupeds [24], whereas McGowan et al. (2008) found that the macropod femur is more robust in larger animals lending support to the concept that intense hopping could relate to increased hindlimb robustness [22]. Musculotendinous forces generated during hopping could incur relatively larger loads on tendon insertion sites around the metaphyses compared to artiodactyls. Those larger loads in macropods may manifest as stronger scaling of cross-sectional parameters in macropods’ metaphyses, evidenced as higher scaling exponents. Conversely, if the more frequently-used, slower gaits’ loading environment drive bone shape then we should expect to see similar scaling between macropods’ fore- and hindlimbs, and between equivalent bones in macropods and artiodactyls, because the low speed pentapedal gait and quadrupedal walking respectively, dominate these clades’ locomotor repertoires.

Using artiodactyls as a quadrupedal comparator clade that has a wide body mass range, we ask whether macropod limb bones exhibit structural scaling features that relate to their pentapedal and hopping locomotor specialisations. In particular, we predict that the forelimb bones of the macropods, which are used for grasping and low-speed locomotion (and are essentially unloaded during hopping), should become relatively more gracile with increases in body size, and consequently have lower scaling exponents than artiodactyl forelimb and macropod hindlimb bones. We hypothesise that scaling exponents should be more similar between fore- and hindlimb bones in artiodactyls than in macropods due to artiodactyls’ more even distribution of stresses between fore- and hindlimbs during high-speed locomotion.

## Materials & Methods

We selected the humerus, radius, ulna, and metacarpal bone (III in macropods and fused III-IV in artiodactyls), along with the femur, tibia and metatarsal bone (IV in macropods and fused III-IV in artiodactyls) from 15 macropod and 20 artiodactyl species (Table 1). All specimens were skeletally mature as determined by fusion of the growth plates, and free from obvious skeletal disease. We imaged the bones in clinical computed tomographic (CT) scanners (LightSpeed 16, Ultra, or Pro 16, GE Medical Systems, Pollards Wood, UK) or for the smallest specimens, in an X-ray microtomographic scanner (XMT; X-Tek HMX ST 225, Nikon Metrology, Tring, UK) with the bone’s long axis positioned parallel to the image’s z- axis, and applied a similar image processing technique used elsewhere [25, 26]. Scans where the long axis of the bone was oblique to the z-axis of the scanner were aligned with BoneJ’s Moments plugin, so that the bone’s principal axes of inertia were parallel with the scan’s x-, y-, and z-axes. Scans with large numbers of image slices were downsampled without interpolation to contain 100-200 slices, providing 5-10 values for averaging in each 5% length bin. Fat in the marrow cavity and other bony or metal elements were manually replaced with a pixel value corresponding to air. Where nearby or fused bones could not be excluded by a rectangular region of interest (ROI), they were manually removed by replacing them with pixels of an air-equivalent value. Bones containing lesions or severe post-mortem deterioration were excluded from the study. Image analysis was performed with BoneJ v1.4.2 [27, 28] for ImageJ v1.51c [29].

**Table 1.**
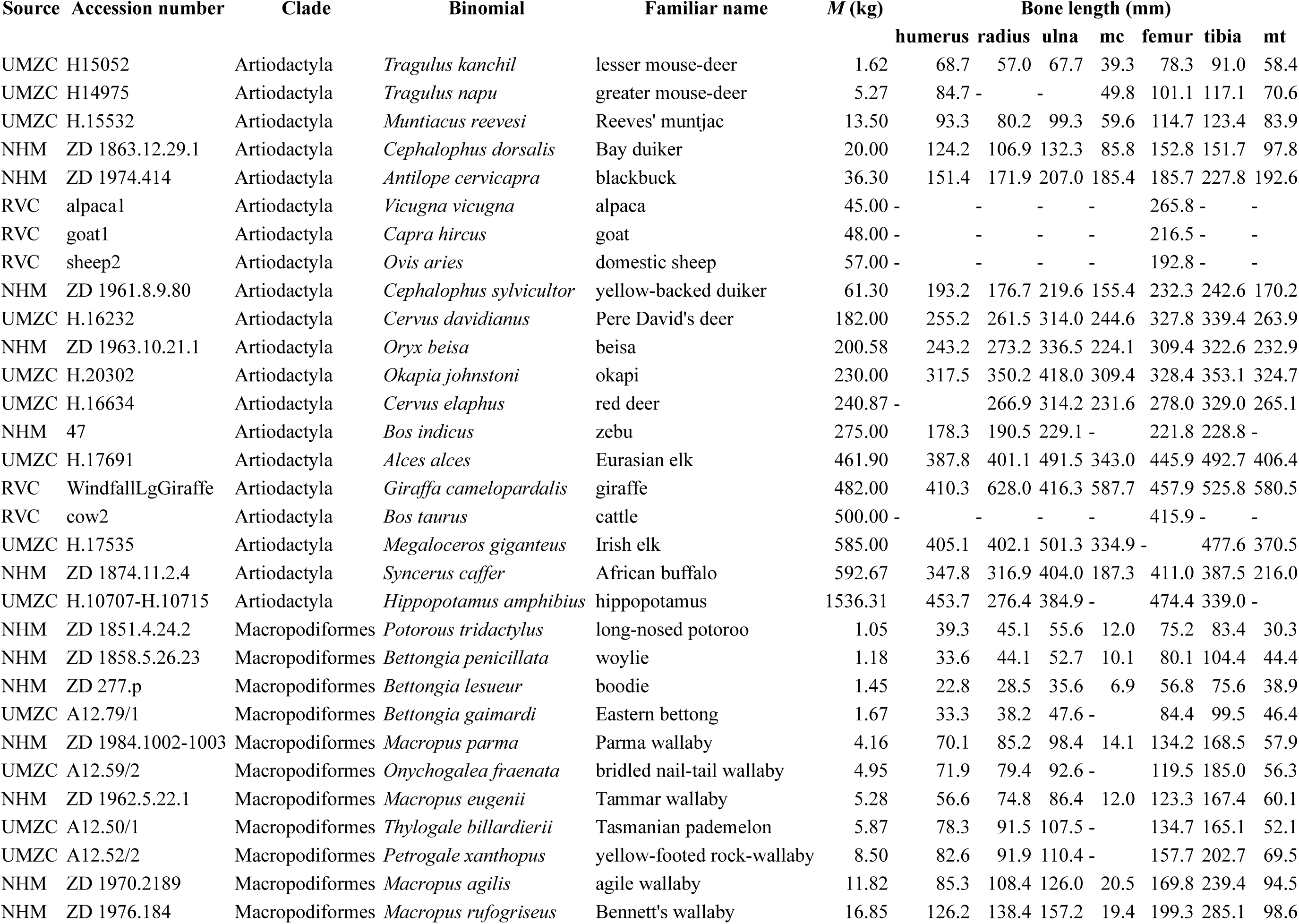
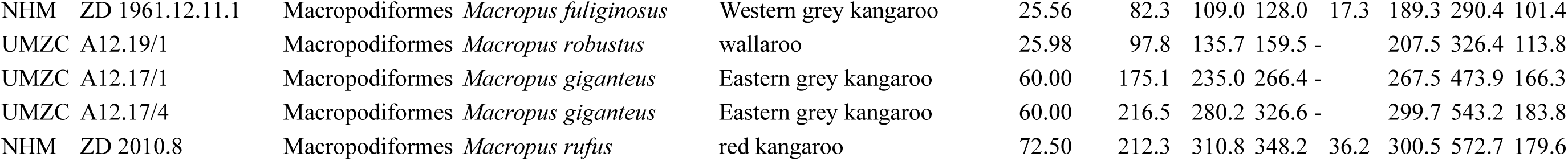
List of specimens. Complete list of specimens, their body masses and lengths of bones used for scaling calculations. Some bones from some specimens were not available to study and are indicated as a dash. NHM, Natural History Museum (London); UMZC, University Museum of Zoology, Cambridge; RVC, The Royal Veterinary College (authors’ collections); mc, metacarpal; mt, metatarsal.

Maximum second moment of area (*I*_max_) was measured on every slice of each scanned specimen with Slice Geometry in BoneJ. Other parameters including *I*_mim_, cross-sectional area and section modulus were also measured and are available in the associated datasets [30], but are not reported here due to their close mathematical relationships: *I* is calculated by multiplying area by distance from the principal axis squared, and section modulus is calculated by dividing *I* by chord length. *I*_max_ can reflect the stiffness of a member in bending, although we use *I*_max_ here as a geometric parameter of cross-sectional size and disposition that includes non-bending related features such as the tibial crest, which loaded mainly in tension by the patellar ligament. To calculate bending stiffness *I* is calculated around the neutral plane of bending, which moves during the stride cycle, and which usually does not coincide with the principal axis used to calculate *I*_max_ [31]. Because the ratios between specimen size, image resolution, and pixel spacing were not constant, we applied a correction for partial filling of pixels which maintains comparable cross-sectional area measurements when image resolution, pixel spacing and resolution vary with respect to each other (Figure 2). Partial filling correction was set by excluding pixels less than −800 HU to eliminate artefacts with values close to air (-1000 HU) and scaling linearly between −1000 HU (0% bone, 100% air) to 2300 HU (100% bone). Pixel values over 2300 HU were considered 100% bone. XMT images lacking HU calibration were set by taking a histogram of an ROI positioned in the background and using its mean for the 100% air scaling value and its maximum as the minimum cutoff value. Another histogram was made in a thick region of cortical bone and its mean used as the 100% bone scaling value. The partial volume correction approach was validated using synthetic images and an exemplar CT image, and resulted in a high degree of stability compared to global thresholding; test scripts and data are available online [30]. Bone length (*l*) was measured using the image data, which we validated against physical measurement of the bones. Body mass (*M*) was unknown for most of the specimens so was estimated from literature values [32–35]. The red and Eastern grey kangaroo specimens were male, so we used body masses near the high end of the estimate to account for the sexual dimorphism in these species.

**Figure 2.**
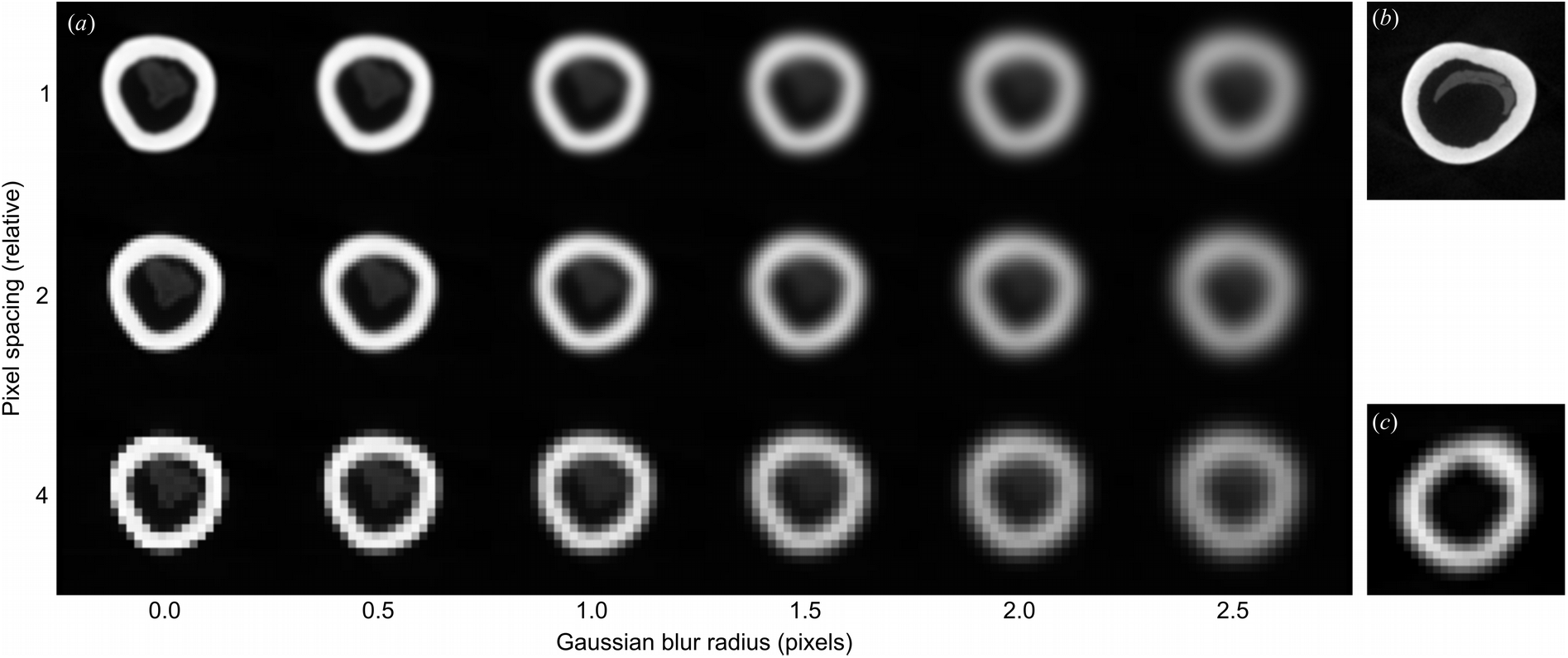
Interaction between specimen size, image resolution and pixel spacing. As pixel spacing increases and resolution decreases relative to specimen size, a greater proportion of pixels represent the edge of the specimens compared to the mid-substance. (a) Progressive downsampling of a well-sampled image of a bone cross-section (top left) increases pixel spacing (vertical axis) and Gaussian blurring with increasing radius simulates lower instrument resolution (horizontal axis). High-resolution images from X-ray microtomography (b) and lower resolution clinical CT images (c) relate to different pixel spacing/image resolution combinations within this scheme. We corrected for imaging condition and specimen size variation using a weighted pixel sum approach in BoneJ’s Slice Geometry plugin.

We analysed scaling of bone dimensions using the general equation *y* = *ax*^b^ [36], where *y* is the bone parameter, *x* is a measure of size (body mass *M* or bone length *l*), *a* is the scaling coefficient and *b* is the scaling exponent. The exponent *b* expresses the rate of change in *y* as a function of body size, while *a* is the magnitude of *y* when *x* = 1. Scaling analysis relies on linear fitting to the log transformed variables, log(*y*) = log(*a*) + *b*log(*x*), where *b* becomes the slope of the line and log(*a*) the *y* intercept or ‘elevation’. The purpose of a scaling analysis is to determine whether relative proportions of the variables under study (here, *l, I*_max_, and *M*) vary with size. Isometry occurs when the larger animal is a ‘to-scale’ version of the smaller animal and the dimensions retain the same proportions, with an isometric scaling exponent *b*_i_. Deviations from isometry occur when *b* is greater or less than *b*_i_, and the corresponding variation in proportions with size is termed allometry. Positive allometry occurs when there is a disproportionately greater increase in *y* with increasing *x* and *b* > *b*_i_, while negative allometry occurs when there is a disproportionately lesser increase in *y* with increasing *x*, and *b* < *b*_i_. Differences in scaling coefficient between groups may be calculated when exponents are equal, and represent a fixed ratio with changing size. Scaling estimates were calculated using smatr version 3.4-3 [37] for R [38], using the standardised major axis (SMA, also known as RMA), which accounts for error in x as well as in y, and is suited to scaling analyses, which typically deal with data that have a large amount of unidentifiable error or “natural variation” [39]. Exclusion of the exponent values for isometry (*l~M*, *b*_i_ = 1/3; *I*_max_~*M*, *b*_i_ = 4/3; *I*_max_~*l*, *b*_i_ = 4) by 95% confidence intervals returned by smatr was used to test the null hypothesis of isometry, and *R*^2^ and *p* from an F-test was used to test the null hypothesis of scale invariance, in which there is no correlation between variables. Cross-sectional parameters were averaged within each 5% increment of length and scaling exponents and elevations calculated for each 5% bin across all the individuals in each clade, for each bone in the study. Normalized cross-sectional parameters were calculated by dividing the *nth* root of the parameter by length. Second moment of area has units of mm^4^, so it was normalized by taking the 4^th^ root and dividing by bone length in mm. Normalized parameters are unitless and a size-independent measure of shape.

To control for non-independence of samples due to their phylogeny, a phylogenetically-informed version of RMA (phyl.RMA) from the R package phytools [40] was called from a custom script to calculate scaling relationships versus M for bone length and *I*_max_ at mid-shaft (50% of bone length). Calibrated phylogenetic trees, one for macropods and one for artiodactyls, were used for this analysis and were constructed based on divergence time estimates from a previous publication [41]; values from the two Eastern grey kangaroo specimens were averaged for the phylogenetic analysis (Figure 3). We report the results of the phylogenetic RMA with five parameters: the estimate of the scaling exponent (b); the squared correlation coefficient (R^2^); the p-value for devation from isometry comparing exponents using an F-test (p); Pagel’s λ, a measure of how well the phylogeny explains the data (λ=1 suggests the evolution of the observed traits follows a Brownian motion model with variance proportional to divergence times, whereas λ=0 suggests phylogenetic independence); and the log-likelihood *(L)* [42] in Table 3. We considered also using phylogenetic generalized least squares (PGLS) analysis, but as prior studies have found negligible differences between phylogenetic RMA and PGLS results or conclusions [43, 44], we opted to apply only phylogenetic RMA to our data, thereby retaining a closer methodological equivalence to our standard scaling analysis.

**Figure 3.**
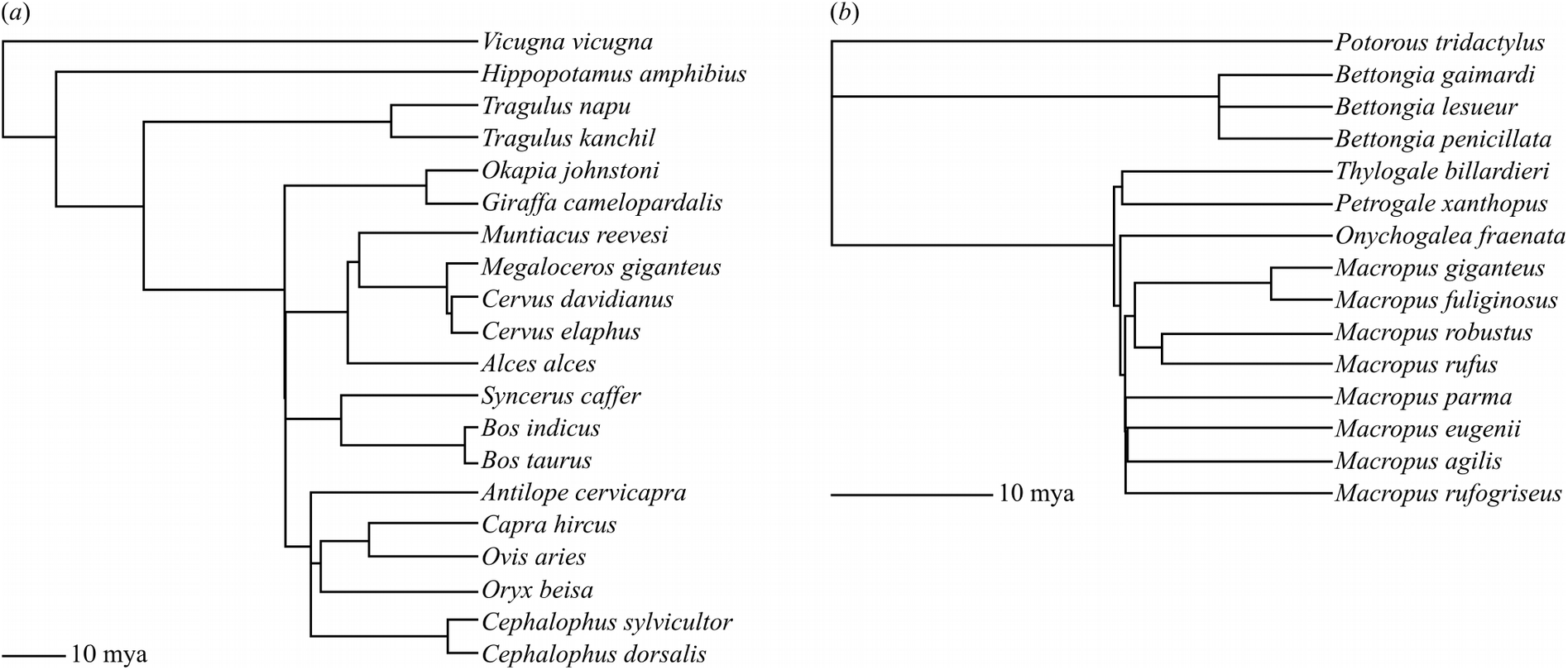
Cladograms illustrating phylogenetic relationships (from [41]) among the artiodactyl (a) and macropod (b) species used to perform phylogenetic independent contrast calculations.

**Table 2.**
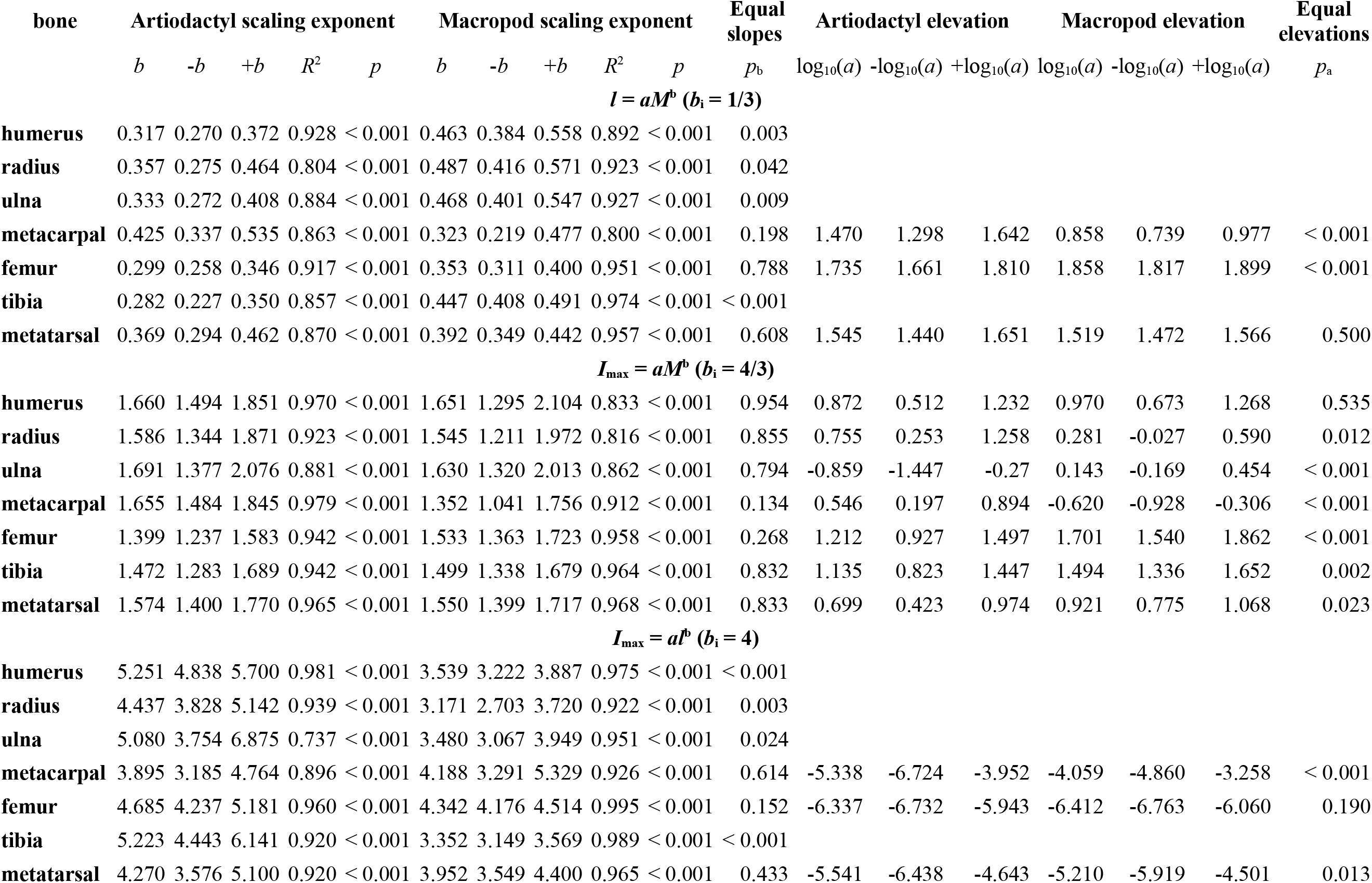
Summary statistics for bone length (*l*) scaling against body mass (*M*), second moment of area (*I*_max_) at 50 per cent length versus *M*; and *I*_max_ at 50 per cent length versus *l* regressions for all bones. The scaling exponent (slope, *b*) is indicated alongside its upper and lower 95% confidence limits (±*b*), along with the coefficient of determination (*R*^2^) and *p* indicating the strength of the correlation between bone length and body mass values (i.e. thep-value of the F-test of the correlation coefficient against zero). Isometric scaling exponents (*b*_i_) are listed for each comparison. The likelihood of equality of scaling exponents between artiodactyl and macropod bones is indicated by the result of a Wald test, *p*_b_. Where slopes are not significantly different, elevations (log_10_(*a*)), their 95% confidence limits (±log_10_(*a*)), and equality of elevations between artiodactyls and macropods (*p*_a_) are reported. Statistical estimates were generated by R using smatr calls in the form: > summary(sma(log10(length)~log10(mass)*order)) > summary(sma(log10(length)~log10(mass)+order, type-’elevation”))

**Table 3.** Bone length and mid-shaft second moment of area phylogenetically-informed RMA scaling exponents against body mass, where *l* « *M*^b^ or *I*_max_ ∝ *M*^b^. The scaling exponent (slope, *b*) is indicated, along with the coefficient of determination (*R*^2^) indicating the strength of the correlation between either *l* or *I*_max_ and body mass values, while *p* denotes thep-value for the isometry test (logarithm of estimated slope is compared to hypothesized value using a t-test, as described in [80]). *λ* denotes Pagel’s lambda, a parameter quantifying phylogenetic signal, while L denotes log likelihood of the fitted model.

| bone | Artiodactyl phylogenetic RMA<br>scaling analysis |  |  |  |  | Macropod phylogenetic RMA<br>scaling analysis |  |  |  |  |
| --- | --- | --- | --- | --- | --- | --- | --- | --- | --- | --- |
| | $b$ | $R^2$ | $p$ | $\lambda$ | $L$ | $b$ | $R^2$ | $p$ | $\lambda$ | $L$ |
| | $l \propto M^b$ | | | | | | | | | |
| humerus | 0.354 | 0.917 | 0.468 | 1.000 | -23.071 | 0.480 | 0.780 | 0.023 | 0.657 | -19.970 |
| radius | 0.401 | 0.855 | 0.108 | 1.000 | -27.833 | 0.489 | 0.843 | 0.005 | 0.562 | -20.015 |
| ulna | 0.373 | 0.880 | 0.270 | 0.982 | -25.363 | 0.471 | 0.853 | 0.007 | 0.553 | -19.015 |
| metacarpal | 0.449 | 0.806 | 0.040 | 1.000 | -27.234 | 0.340 | 0.726 | 0.919 | 0.514 | -13.808 |
| femur | 0.314 | 0.894 | 0.455 | 0.548 | -27.826 | 0.346 | 0.894 | 0.695 | 0.592 | -11.671 |
| tibia | 0.330 | 0.858 | 0.917 | 1.000 | -25.230 | 0.446 | 0.949 | 0.001 | 0.409 | -11.396 |
| metatarsal | 0.401 | 0.824 | 0.158 | 1.000 | -24.973 | 0.432 | 0.935 | 0.004 | 0.818 | -9.837 |
| | $I_{max} \propto M^b$ | | | | | | | | | |
| humerus | 1.740 | 0.951 | 0.001 | 0.842 | -43.370 | 1.726 | 0.640 | 0.164 | 0.669 | -41.410 |
| radius | 1.688 | 0.909 | 0.017 | 0.827 | -47.003 | 1.609 | 0.645 | 0.279 | 0.539 | -44.176 |
| ulna | 1.777 | 0.914 | 0.005 | 0.976 | -46.334 | 1.668 | 0.717 | 0.156 | 0.587 | -42.659 |
| metacarpal | 1.725 | 0.956 | 0.002 | 0.675 | -37.438 | 1.409 | 0.880 | 0.685 | 0.429 | -23.174 |
| femur | 1.399 | 0.942 | 0.424 | 0.000 | -54.017 | 1.496 | 0.908 | 0.199 | 0.591 | -32.535 |
| tibia | 1.552 | 0.914 | 0.078 | 0.697 | -48.396 | 1.469 | 0.927 | 0.243 | 0.419 | -30.572 |
| metatarsal | 1.651 | 0.950 | 0.008 | 0.633 | -37.981 | 1.564 | 0.941 | 0.038 | 0.407 | -31.343 |

## Results

Bone length versus body mass, and 50 per cent length *I*_max_ versus *l* and *M*, comparisons are presented in Table 2 and Figure 4. *I*_max_, *l*, and *M* are strongly correlated in all bones with high *R*^2^ (0.737-0.995) and *p* < 0.001, excluding the null hypothesis of no scaling relationship between the variables. Humerus, radius and ulna lengths scale with positive allometry (*b* > 1/3) in macropods, but with isometry (b not significantly different from 1/3) in artiodactyls. Artiodactyl metacarpal bones are much longer than in macropods of similar mass, indicated by the high elevation (1.47 vs. 0.86). In the hindlimb, femur and metatarsal lengths scale similarly in macropods and artiodactyls, with the macropod femur having a higher elevation than artiodactyls and the metatarsals’ slopes and elevations not significantly different. Tibia length scales isometrically in artiodactyls and with strong positive allometry in macropods.

**Figure 4.**
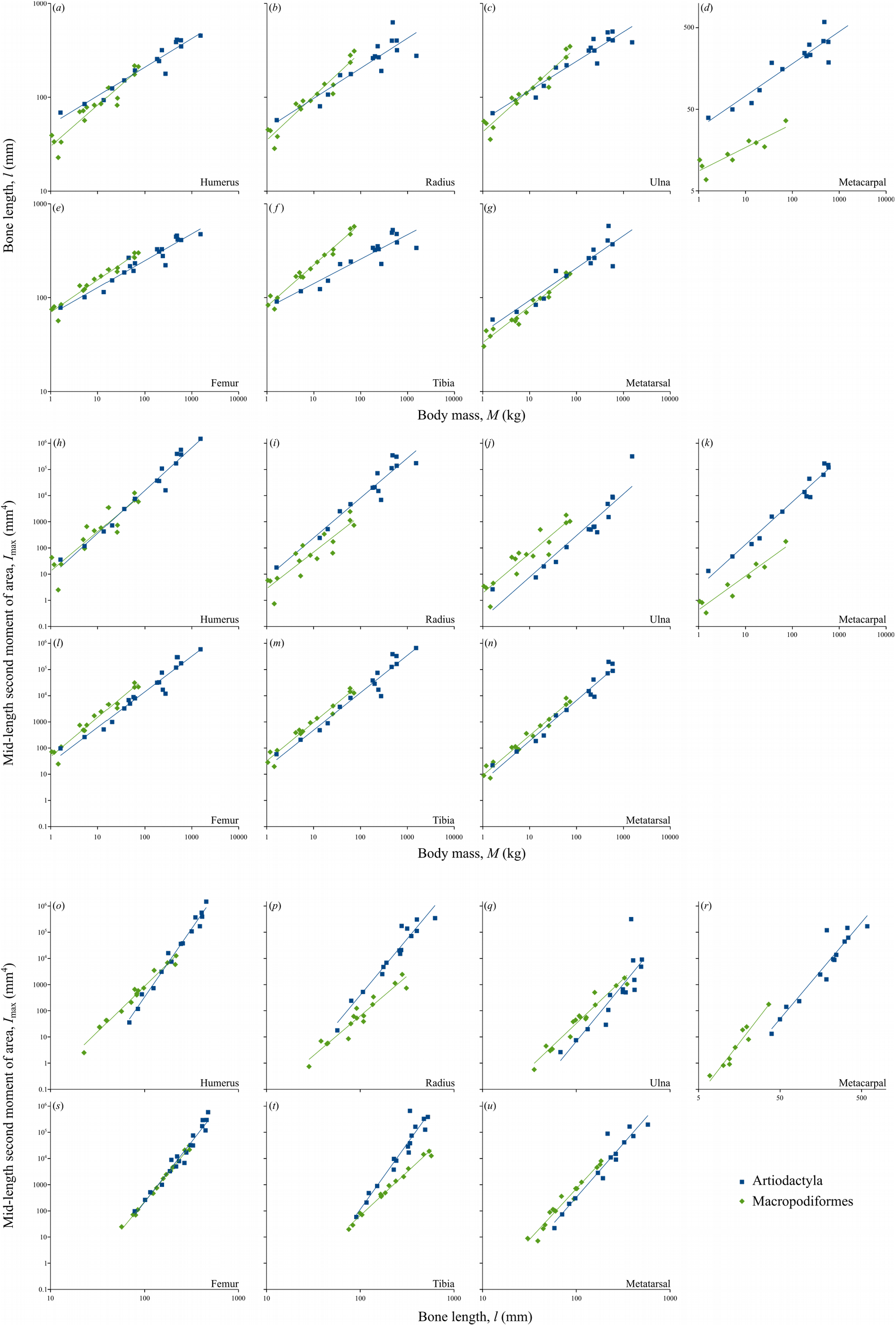
Bone length (*l*) versus body mass (*M; a-g*);second moment of area (*I*_max_) at 50 per cent length versus *M*(*h-n*); and *I*_max_ at 50 per cent length versus *l*(*o-u*) regressions for all bones. Bone lengths and body masses are presented in Table 1. Scaling exponents (slopes, *b*), elevations (log10(a)), *R*^2^ and *p* values are presented in Table 2.

Comparing stylopod (humerus, femur), zeugopod (radius, ulna, tibia), and autopod (metacarpal, metatarsal) elements between limbs within each of the two clades, there is a high degree of overlap between the confidence limits of scaling exponents in all the limb segments, meaning that bone length proportionality between fore- and hindlimb segments is maintained within clades. Mid-diaphyseal *I*_max_ scales against *M* with similar exponents between clades but with differing elevations in all bones except the humerus, indicating constant proportionality between clades with increasing animal size. Mid-diaphyseal *I*_max_ scales against *l* with different exponents between clades in the humerus, radius, ulna and tibia, and with different elevations in the metacarpal and metatarsal. Notably, the femora are indistinguishable at 50 per cent length in their *I*_max_ ~ *l* scaling.

Normalized *I*_max_ versus per cent length plots (Figure 5) reveal that artiodactyls’ cross-sections become relatively more robust with increasing body mass, indicated by the larger animals’ traces tending towards the top of the range. Meanwhile, macropods show the opposite trend, with normalized *I*_max_ decreasing with increasing body mass so that traces from the larger animals appear at the bottom of the range, indicating increased gracility with increasing body mass. In general, and in common with prior studies on cats and birds [25, 26], the artiodactyl diaphysis occupies a decreasing proportion (and metaphyses and epiphyses increasing proportions) of bone length with increasing body mass, but this relationship is not maintained consistently in macropods. Notably, the trochlear notch and coronoid processes of the ulna drift distally in larger artiodactyls, but proximally in larger macropods (Figure 5e, f).

**Figure 5.**
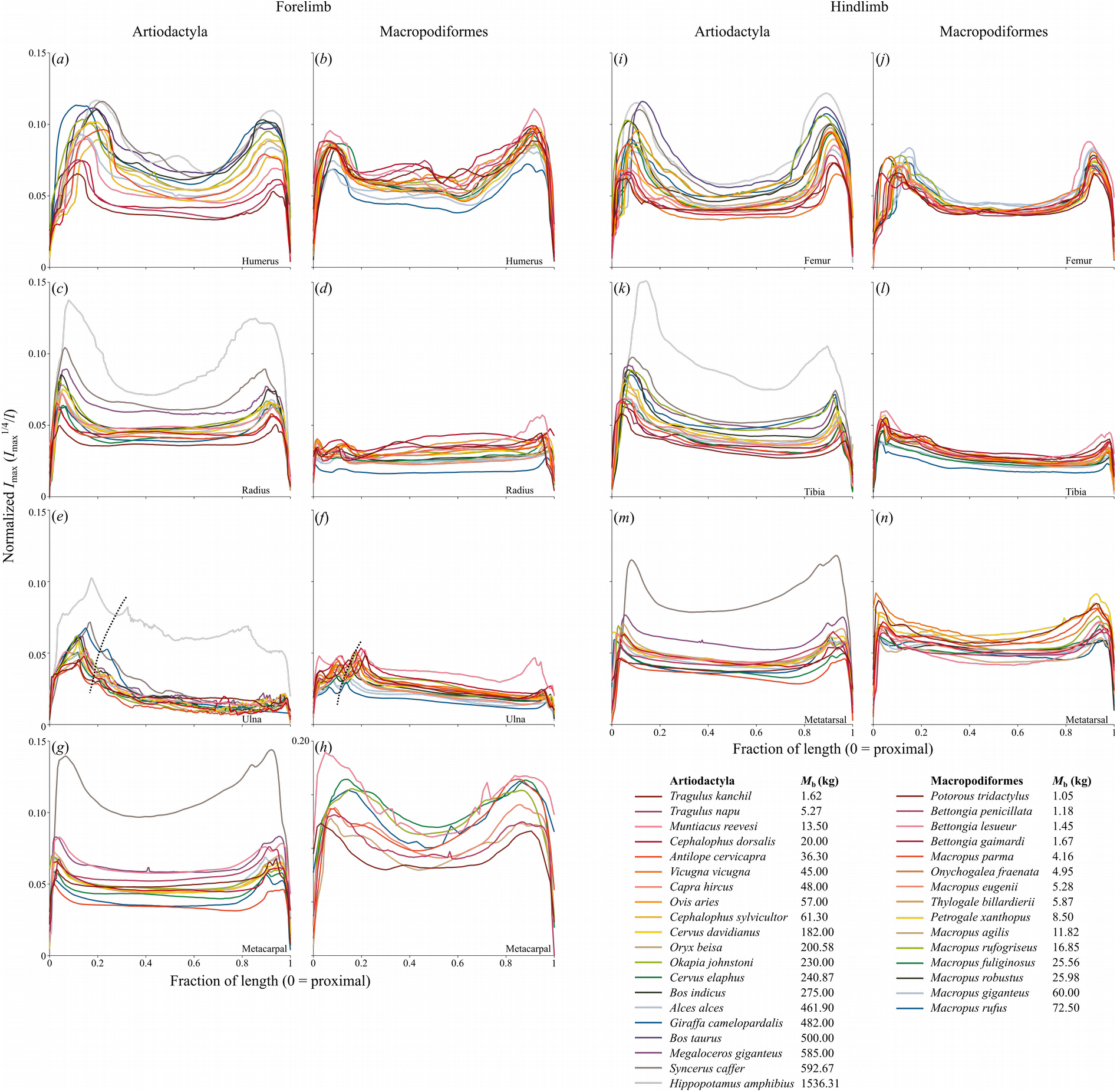
Normalized second moment of area (*I*_max_^1/4^/*l*) at each fraction of length. Dispersal of traces indicates bone shape differences among species. Higher traces indicate relatively more robust bone geometry, seen in larger artiodactyls and smaller macropods. Note the increasing proportion of length occupied by epiphyseal and metaphyseal components in larger artiodactyl species (*a, i*), and the distal drift of the ulna’s trochlear notch (dashed line) in larger artiodactyls (*e*) and smaller macropods (*f*).

Scaling exponents (Figure 6) and elevations (Figure 7) for *I*_max_ versus *M* reveal near-identical scaling exponents between clades for all regions of all the bones, and overlapping elevations for all bones in all regions except for the proximal tibial and femoral metaphyses, indicating very similar bone cross-sectional scaling against body mass. Positive allometry (exponent above the isometry line) is strongest in the proximal metaphyses, possibly relating to their increasing relative length mentioned above, and this is amplified by increased elevations (i.e. larger value of *I*_max_ at a given *M*) in these regions in macropods (Fig 7f, l, n). *I*_max_ versus *l* scaling reveals positive allometry for much of the length of artiodactyl bones. The wide confidence interval of artiodactyl ulna (Figure 6e) likely reflects the variability of fusion to the radius, reducing the strength of the body size signal. In contrast, macropod *I*_max_ scales with negative allometry against *l* for much of the length of humerus, radius, ulna and tibia, with positive allometry in the femur and isometry in the metacarpal and metatarsal. The raised elevation of macropods relative to artiodactyls in the *I*_max_ versus *l* plots (Figure 7) is difficult to interpret because the scaling exponents are markedly different between clades in the regions where elevations are different. Despite their orders of magnitude difference the elevations may not relate to functional differences, which may be more strongly indicated by differing scaling exponents.

**Figure 6.**
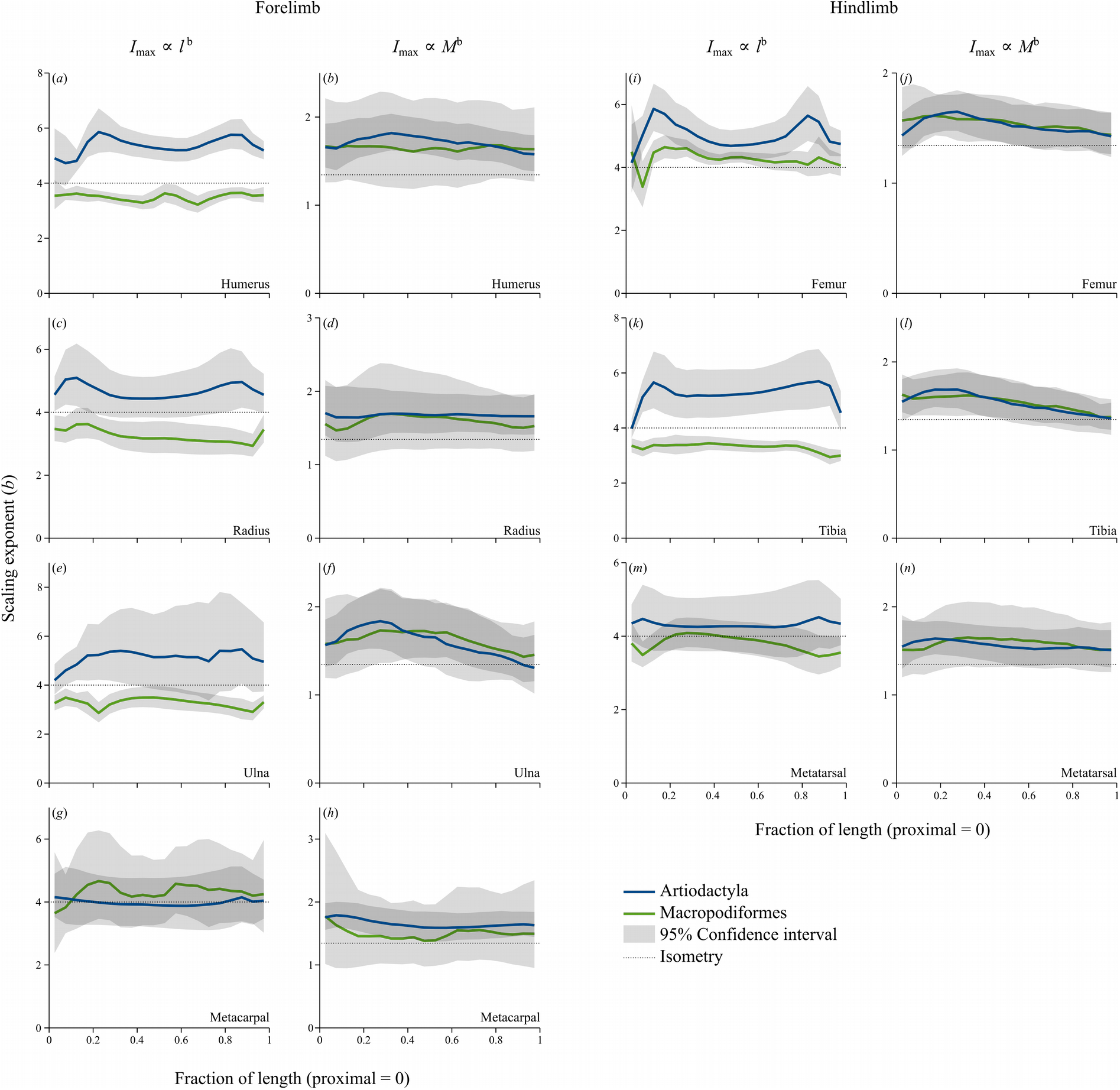
Scaling exponents for second moment of area (*I*_max_) versus bone length (*l*) and body mass (*M*) for all bones and both clades. Light grey regions indicate the 95% confidence interval; dark grey regions occur where the confidence intervals overlap and where substantial may be interpreted as no significant difference in scaling exponent in that region of the bone, between clades. Scaling exponent estimates at 50 per cent length are presented in Table 2.

**Figure 7.**
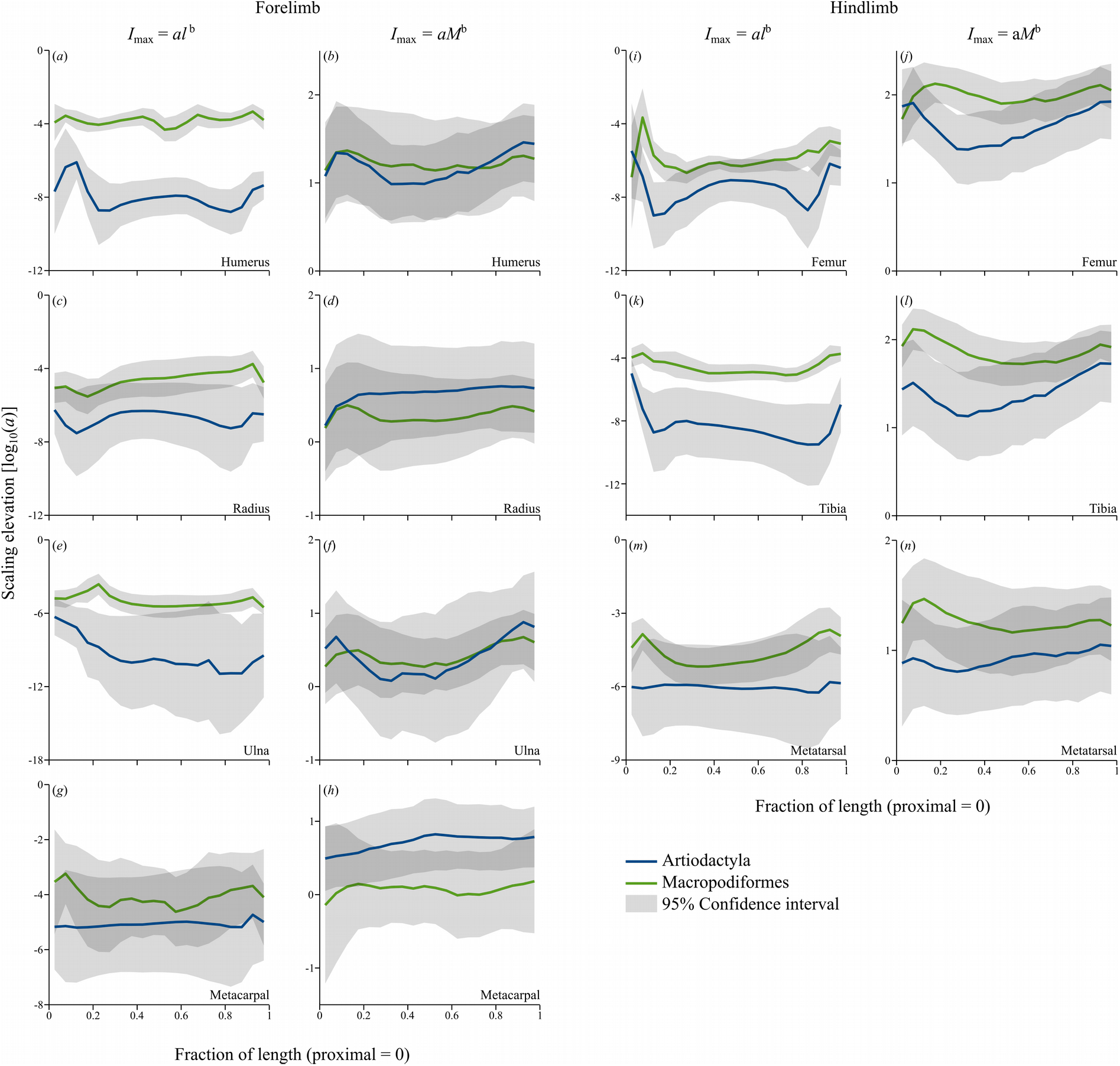
Scaling elevations [log_10_(*a*)] for second moment of area (*I*_max_) versus bone length (*l*) and body mass (*M*) for all bones and both clades. Elevations are directly comparable only where slopes (scaling exponents, Figure 6) are equal. Light grey regions indicate the 95% confidence interval; dark grey regions occur where the confidence intervals overlap and where substantial may be interpreted as no significant difference in scaling elevation between clades. Scaling elevation estimates at 50 per cent length are presented in Table 2.

Despite there being a highly variable amount of phylogenetic signal (Pagel’s λ ranged between 0.407 and 0.818 for macropod bones, while it covered the entire range of 0-1 for artiodactyl bones), scaling exponents for both artiodactyl and macropod bone length and midshaft *I*_max_ corrected for phylogenetic effects using phylogenetically-informed RMA remained comfortably within the 95% confidence limits of the scaling exponents calculated without phylogenetic adjustment (Tables 2 and 3). Correlations calculated using phylogenetic RMA were similarly strong as their non-corrected counterparts (unadjusted *R*^2^ = 0.800-0.979, adjusted *R*^2^ = 0.640-0.956).

In our phylogenetically-informed RMA, macropod femur and third metacarpal lengths scale isometrically, while all other macropod bone lengths scale with positive allometry. Macropod mid-shaft *I*_max_ does not scale differently from isometry in all bones except the metatarsal (*p*: humerus 0.164; radius 0.279; ulna 0.156; metacarpal 0.685; femur 0.199; tibia 0.243; metatarsal 0.038).

Phylogenetically-informed RMA suggests that artiodactyl bone lengths scale isometrically except for the metacarpal (slope not different from 1/3,p: humerus 0.468; radius 0.108; ulna 0.27; metacarpal 0.04; femur 0.455; tibia 0.917; metatarsal 0.158). Artiodactyl mid-shaft *I*_max_ scales with positive allometry in the humerus, ulna, metatarsal and metacarpal (p<0.005), with a similar tendency in the radius (p=0.017), but not the femur or tibia (*p*: femur 0.424; tibia 0.078).

In summary, scaling exponents calculated using phylogenetic adjustments for bone length, mid-shaft CSA and mid-shaft *I*_max_ concur with the results from the unadjusted scaling relationships. Thus phylogeny has varying explanatory value for bone geometry scaling (as quantified by Pagel’s λ) within the two clades of mammals studied here, but does not influence the interpretation of the scaling exponents, as the estimates are essentially no different between phylogenetically informed and non-phylogenetic scaling analyses.

## Discussion

Scaling of the forelimb and hind limb segments is similar within clades, except the stylopod, in which the *I*_max_ versus length scaling displays positive allometry in the artiodactyl and macropod femur and artiodactyl humerus, but negative allometry in the macropod humerus, meaning that in macropods the humerus becomes more gracile with increasing length while the femur becomes more robust. Unlike Bennett (2000), who found that tibial second moment of area scales more strongly positively in kangaroos than quadrupeds (b = 1.52 *vs* 1.28) [24], our data show that tibial cross-sections scale similarly against body mass between clades. This may be a consequence of comparing macropods to artiodactyls only, and not to a more diverse sample of quadrupeds, because it is known that artiodactyls’ bones scale differently to other mammalian clades [19, 20]. Tibial cross-sections scale strongly negatively allometrically in macropods and positively allometrically in artiodactyls against length. This means larger kangaroos’ tibiae are relatively less robust – they are relatively longer and more slender consistent with a relatively reduced ability to resist bending moments. This apparent reduction in relative bending strength is surprising considering that bending stresses predominate over compressive stresses due to the off-axis component of the muscular forces, with a stress range of −110 to −60 MPa and 90-110MPa [1].

The *I*_max_ versus body mass scaling elevation is higher in the macropod proximal femur and tibia than the same region in the artiodactyl femur and tibia, indicating increased robustness around the greater and lesser trochanters and tibial crest, which are the bony insertions for the massive gluteal, iliopsoas, and quadriceps muscle groups that drive bipedal hopping in macropods. Positive allometry of tarsal joint moment arms potentially ameliorates the musculotendinous compressive force on the tibia during tarsus extension [22], allowing the distal half of macropods’ tibial cross-sections to remain within similar parameters as artiodactyls’. Like McGowan et al. (2008) demonstrated in macropods, we find that the femur is more robust in larger macropods [22], which is consistent with a proposal of a strong, general relationship between stylopod cross-sectional parameters and body mass [21]. We find that humeral and femoral lengths scale significantly differently against body mass between macropods and artiodactyls, in contrast to suggestions of common mammalian femur length to body mass scaling [45], which may have implications for midshaft bending stresses.

The largest extant artiodactyls are an order of magnitude more massive than the largest extant macropods while the smallest of both clades included in this study are ~1-2 kg. It would be unwise to extrapolate macropod scaling trends beyond the current series, because bipedal hopping was likely not a feature of the extinct giant kangaroos and may not be physiologically possible beyond ~160 kg [7–10]. Janis et al. (2014) suggested that large, extant kangaroos are functionally specialised for hopping in contrast to their larger extinct kin that did not hop, somewhat similar to the medium-sized, gracile and hyper-athletic cheetah (*Acinonyx jubatus, M* = 35-70 kg) compared to bigger and more robust felids such as lion (*Panthera leo, M* = 120-250 kg) [9].

We found that the trochlear notch of the ulna is relatively more distal in larger artiodactyls, but that an opposite trend of a relatively shortened olecranon process and proximally drifting trochlear notch is observed with increasing mass in macropods. We first noticed a trend to a more centrally-placed trochlear notch in the ulna of large felids [25], and proposed that this may be a mechanism that may allow reduced muscle forces by increasing the lever arm of the olecranon process and increasing the elbow extensor muscles’ effective mechanical advantage. The relatively shortening olecranon in larger macropods may relate to forelimb use in the low-intensity pentapedal gait and lack of loading in high-intensity bipedal hopping, or to reaching and combat behaviours favouring a longer forelimb. Inter-clade differences in metacarpal dimensions relate to their functional specialisations for grasping, or plantigrade or unguligrade locomotion in macropods and artiodactyls respectively. Similar isometric *I*_max_ scaling exponents against length indicate maintenance of overall bone shape that may relate to specialised manus function, whereas positive allometry against body mass in artiodactyls but isometry in macropods may reflect an influence of locomotor loading on artiodactyl metacarpal robustness that is absent or reduced in macropods.

Bones respond anabolically, that is, by increasing bone tissue formation and decreasing bone resorption, when they experience a small number of novel high strain and high strain rate events with a rest period between bouts of loading [46, 47]. Repetitive loading has a saturation or habituation effect, in which tissue is no longer responsive to mechanical loads after a few tens of cycles [47]. The lack of a difference in femoral and tibial *I*_max_ versus body mass scaling exponents between bipedal hopping macropods and quadrupedal artiodactyls suggests that the occasional very high load of hopping may not be sufficient to overcome the mechanobiological saturation engendered by frequent but lower intensity loading in crouching and pentapedal walking. Alternatively, bipedal hopping may be a no more intense stimulus to the hindlimb than quadrupedal galloping, leading to little discernible difference between clades. Galloping and hopping engender similar muscle stresses at preferred speeds in rats and kangaroo rats respectively despite a fourfold difference in ground reaction force [67]. We found similar *I*_max_ ~ *M* scaling exponents in the forelimb bones, despite macropods’ relatively unloaded forelimbs during bipedal hopping.

Limb bone adaptation occurs throughout the life of the individual due to bone’s phenotypic plasticity, and at the population level over evolution due to selection pressure relating to skeletal development and functional specialisation. Variable safety factors among species and bones [48–50] suggest that peak strains from uncommon or high energy gaits are not necessarily the dominant stimuli for bones’ phenotypic adaptation, or a critical selection pressure in evolution. Bone is sensitive to its mechanical environment during growth: altered muscle and gait forces on growing bones relate to bone deformities in children with cerebral palsy and in experimentally manipulated chick and crocodile embryos [51, 52], while 12 hours per week of throwing practice leads to substantial cortical bone adaptation in children’s throwing arms [53]. In-vivo bone adaptation experiments have shown that the mouse tibia receives < 300 με (microstrain) during walking and < 600 με when landing a 30cm jump (~ 3N physiological load), yet requires at least 1000 με from a 10N experimental load to stimulate further bone formation [54–56]. Only high intensity race training, well beyond the animals’ *ad libitum* locomotor behaviour, was sufficient to engender an increase in cortical area in horses [57]. Sciatic neurectomy removes daily habitual loading in the mouse hindlimb, sensitising the tibia to subsequent load-induced (2000 με) bone deposition [58], suggesting that the removal of background stimulus can rescue bone’s load responsiveness [59], or in other words, that daily stimulus might saturate bone’s ability to respond to further applied loads. Simple body mass support and the ground reaction forces incurred by a slow gait may be sufficient to maintain cross-sectional bone geometry, while infrequent high intensity quadrupedal gaits might offer little further selection pressure or modelling stimulus to increase diaphyseal size over and above that provided by standing and walking.

There are few data on the daily numbers of stride cycles in each gait for the species in the study, which limits our ability to calculate bone loading histories and infer which gaits relate most strongly to bone structural scaling, however, in those species that have been studied low intensity behaviours predominate. In large macropods, the most frequent behaviour is lying down or standing still, followed by slow locomotion and only very occasional hopping [3, 5]. Basic data on macropod locomotor activity exist in addition to that of the tammar wallaby already mentioned [6]. Locomotion comprises only 5-10% of the behavioural repertoire of the parma wallaby *(Macropus parma)* [60]. The larger red and grey kangaroos *(Macropus rufus* and *M. giganteus)* spend the day alternating between lying, standing, crouching, grazing, and licking [5]. Agile wallabies’ *(Macropus agilis)* most common behaviour is foraging (73%), followed by ‘vigilance’ (23%), and locomotion (0-6%) [61]. In grey kangaroos, over 90% of daily activity is crouching and lying, with only 0.0-3.3% accounted for by ‘moving’ [62]. Artiodactyls are similarly slow most of the time: wildebeest *(Connochaetes* spp.) travel only 2-3km daily [63]; red deer *(Cervus elaphus)* move on average 100-400m per hour [64], while giraffe may walk for 5h daily [65] and can canter for only a few minutes at a time [66].

A scaling trend in gait preference might exist, such that small animals hop, trot or gallop more frequently than large animals, which could influence the interpretation of our results. Macropod species that live in open country generally have a shorter period of suspension than those that live in dense forests or rocky hills, with a potential phylogenetic contribution to duty factor [2] and thus peak ground reaction forces and bone strains. Our phylogenetically-corrected scaling analysis found only limited effects of phylogeny on skeletal scaling parameters, suggesting little relationship between behavioural ecology, locomotor style, and bone geometry scaling within macropod and artiodactyl clades. Comprehensive behavioural ecology, activity pattern, kinematic, ground reaction force, and tissue strain data would help to place the skeletal scaling that we have identified into the context of functional loading. Examining the skeletal scaling of closely related quadrupeds from the diprotodont order such as wombats (Vombatidae), koala (*Phascolarctos*), and possums (Phalangeridae) in the context of hopping macropods’ skeletal scaling, may help to further separate phylogenetic from mechanobiological effects. Kinematic data exist for sheep, goats [68, 69], pigs [70], cattle and a small number of other artiodactyls during walking [71] and for a small number of macropods [72–74], but bone strain data are missing in all but a few species [69, 75].

The lack of differential cross-sectional scaling in the macropod hindlimb despite their hopping behaviour led us to the speculation that their bones might have enhanced fatigue damage repair by increased remodelling, thus reducing the need for extra bone mass. We failed to find secondary osteonal remodelling in a *Macropus giganteus* femur sample, which was somewhat unusual for an animal of 33kg body mass [76]. Absence of secondary osteons may relate to the single sample failing to include any by chance, a load-related suppression of remodelling protecting bone from local weakening due to osteoclastic resorption [57], or infilling of existing osteons as occurs in horses after moderate-intensity training [77]. The current and other studies of bone organ allometry assume no size-related variation in bone microstructure or physiology and that all mammalian bone has similar biomechanical and mechanobiological behavior. Our recent work demonstrated that secondary osteons are wider in larger animals and narrower in smaller animals [76], and that trabeculae are thicker and more widely spaced in larger animals [78] indicating that biophysical constraints or cellular behaviour may vary among mammals and potentially interact with whole-bone-level scaling. Integration of macro- and micro-level perspectives in future scaling studies could be particularly informative.

Forelimb-hindlimb and bipedal-quadrupedal comparisons of scaling relationships have revealed very similar cross-sectional scaling against mass in the primary weightbearing limb bones in artiodactyls and macropods, despite differences in their high intensity gaits, suggesting that habitual low loads rather than occasional high loads may be the dominant stimuli for bone modelling (i.e. scaling) in individuals and across evolution. Cross-sectional scaling against length meanwhile appears to relate to clade-related specialisations such as macropods’ long, gracile forelimb used in low-speed weightbearing and grasping food, and artiodactyls’ more robust forelimb bones specialised for cursorial locomotion.

## Data Accessibility

Code, scripts, and databases [30] and raw and processed images [79] are available on figshare under a CC-BY licence. BoneJ is available from bonej.org and v1.4.2 source code is at zenodo [28].

## Authors’ contributions

MD collected and imaged specimens, wrote code, analysed the data, and drafted the manuscript; AAF performed the phylogenetic scaling analysis and helped to draft the manuscript; MMK performed X-ray microtomography; MYC and KL performed preliminary analyses; SJS and JRH conceived of and designed the study, and helped draft the manuscript. All authors gave final approval for publication.

## Acknowledgements

For help with specimen loans we thank Matt Lowe at the University Museum of Zoology, Cambridge, and Roberto Portela-Miguez and Louise Tomsett at the Natural History Museum London. Richard Abel assisted with X-ray microtomography, Renate Weller and Charlotte Mumby assisted with computed tomography, Andrew Cuff advised on phylogenetic scaling analysis and Alexis Wiktorowicz-Conroy gave valuable advice regarding macropod gait. We thank Andrew Pitsillides and Behzad Javaheri for helpful conversations about bone mechanobiology, three reviewers, and the editors for their comments that have helped us to improve the manuscript.

